# Phosphorylation-dependent interactions of VAPB and ELYS contribute to the temporal progression of mitosis

**DOI:** 10.1101/2023.07.03.547506

**Authors:** Christina James, Ulrike Möller, Sabine König, Henning Urlaub, Ralph H. Kehlenbach

## Abstract

ELYS is a nucleoporin that localizes to the nuclear side of the nuclear envelope in interphase cells. In mitosis, it serves as an assembly platform that interacts with chromatin and then with nucleoporin subcomplexes to initiate the formation of novel nuclear pore complexes. Here we describe the interaction of ELYS with the membrane protein VAPB. In mitosis, ELYS becomes phosphorylated at many sites, including a predicted FFAT (two phenylalanines in an acidic tract) motif, which is shown to mediate interaction with the MSP (major sperm protein)-domain of VAPB. Phosphorylation-dependent binding of VAPB to ELYS is demonstrated by peptide binding assays and co-immunoprecipitation experiments. In anaphase, the two proteins co-localize to the non-core region of the newly forming nuclear envelope. Depletion of VAPB resulted in prolonged mitosis and slow progression from meta-to anaphase and also to chromosome segregation defects. Together, our results suggest an active role of VAPB in recruiting membrane fragments to chromatin and in the biogenesis of a novel nuclear envelope during mitosis.

## Introduction

The Vesicle-Associated-membrane Protein-associated protein B (VAPB) is a tail-anchored membrane protein with well-described functions at the endoplasmic reticulum (ER). It serves as a tethering factor that is involved in the formation of contact sites between the ER and other organelles, e.g. mitochondria (De Vos et al., 2012; Gómez-Suaga et al., 2019; Stoica et al., 2014), the Golgi-apparatus (Kuijpers et al., 2013) or peroxisomes (Costello et al., 2017; Kors et al., 2022). Prominent examples for protein-protein interaction partners are VAPB and the oxysterol binding proteins (OSBPs), connecting ER- and Golgi-membranes (Mesmin et al., 2013) and VAPB and the acyl-CoA binding domain protein ACBD5, forming a link between the ER and peroxisomes (Costello et al., 2017). Together, more than 250 interaction- or proximity partners of VAPB have been identified in proteomic screens (Huttlin et al., 2015; James and Kehlenbach, 2021; James et al., 2019; Murphy and Levine, 2016; Slee and Levine, 2019). The biological significance of most of these interactions, however, has not been investigated in detail. Besides its characteristic C-terminal transmembrane domain, VAPB contains an N-terminal MSP (Major Sperm Protein) domain and a central coiled-coil domain (Nishimura et al., 1999). With respect to protein-protein interactions, the MSP-domain is of particular importance. The crystal structure of an MSP-domain of VAPA, a protein closely related to VAPB, revealed seven immunoglobulin-like β-sheets (Kaiser et al., 2005; Shi et al., 2010). The domain interacts with characteristic peptide sequences of target proteins called FFAT-motifs (two phenylalanines in an acidic stretch), which are conserved in OSBPs and in OSBP-related proteins (ORPs). Several variants of such motifs have been described: (i) the conventional FFAT motif with its seven core residues (EFFDAxE) has an upstream acidic flanking region and binds in a characteristic manner to the MSP-domain, where two amino acid residues are crucial for the interaction (VAPB K87 and M89). Hence, exchanging these amino acids to aspartic acid largely abolishes the interaction of VAPB to FFAT-dependent target proteins. (ii) FFAT-like motifs come in different flavors and may even lack the two phenylalanine residues (Murphy and Levine, 2016). They may exhibit a certain binding preference to specific members of the VAP (VAMP-associated proteins; VAMP stands for vesicle-associated membrane protein) family of proteins. Besides VAPA and VAPB, the Motile Sperm Domain-containing proteins 1, 2 and 3 (MOSPD1, MOSPD2 and MOSPD3 (Cabukusta et al., 2020; Di Mattia et al., 2018; Loewen and Levine, 2005)) and the Cilia and Flagella-Associated Protein 65 (CFAP65; (Tang et al., 2017)) belong to this family, all of which contain the characteristic MSP-domain. Both, the canonical FFAT- and the FFAT-like motifs can be regulated by phosphorylation. The negative charge that can be introduced by phosphorylation of serine or threonine residues can promote binding of the respective protein to the MSP-domain. Examples for proteins with such a motif are STARD3 (StAR-related lipid transfer domain-3), a protein that mediates cholesterol transport from the ER to endosomes (Di Mattia et al., 2020), PTPIP51 (protein tyrosine phosphatase interacting protein-51), which tethers mitochondria to the ER (De Vos et al., 2012) and CERT (ceramide transfer protein), which is required for the transfer of ceramide from the ER to the Golgi apparatus (Kumagai et al., 2014). For a more detailed discussion of the different types of FFAT-motifs see (Di Mattia et al., 2020; James and Kehlenbach, 2021; Murphy and Levine, 2016; Slee and Levine, 2019).

Using functional assays and immunoelectron microscopy, we (James et al., 2019) and others (Saiz-Ros et al., 2019) recently showed that VAPB not only localizes to the ER, but also reaches the inner nuclear membrane (INM). Proteins with a cytoplasmic region smaller than 60 kDa are thought to passively diffuse from the ER via the outer nuclear membrane (ONM) and the nuclear pore complexes (NPCs) to the INM (Zuleger et al., 2012). Hence, a portion of the cellular VAPB with its cytoplasmic domain of 24 kDa is expected to reach the INM and the question arises whether the protein fulfills a function at this localization. To address this question, we searched specifically for nuclear interaction/proximity partners of VAPB using an APEX2-based approach for the identification of potential binding partners. Our new method, RAPIDS (Rapamycin and APEX dependent Identification of proteins by SILAC), identified several membrane proteins of the INM, e.g. emerin and LAP1 (lamina-associated polypeptide 1, also known as Torsin-1A-interacting protein 1, TOR1AIP1). Another prominent hit we obtained was the nucleoporin ELYS (embryonic large molecule derived from yolk sac), also known as AHCTF1 (AT-hook containing transcription factor 1) and Mel28 in *C. elegans* (Galy et al., 2006). It was originally identified in mice as a large protein of 2243 amino acid residues (2266 in human cells), containing characteristic nuclear localization signals (NLSs) and nuclear export signals (NESs). Furthermore, an AT-hook DNA-binding domain could contribute to the nuclear localization of the protein (Kimura et al., 2002). Later, it was realized that ELYS is a component of the nuclear pore complex (NPC) and also localizes to kinetochores during mitosis (Rasala et al., 2006). At the level of the NPC, it interacts with the Nup107-160 complex, also known as the Y-complex (Galy et al., 2006; Rasala et al., 2006). Very detailed analyses of NPC-structures revealed that ELYS localizes to the nuclear side of the nuclear envelope (NE) (Bley et al., 2022). Importantly, ELYS is required for the assembly of novel NPCs during mitosis (Franz et al., 2007; Galy et al., 2006; Gillespie et al., 2007; Rasala et al., 2006). Specifically, it binds to the decondensing chromosomes via its AT-hook and then recruits other NPC-components, e.g. other proteins of the Y-complex. This initial hub was then suggested to interact with membrane vesicles containing the integral NPC-membrane proteins POM121 and NDC1 (Rasala et al., 2008), followed by the recruitment of additional soluble nucleoporins. Additional membrane proteins on mitotic membrane fragments that interact with ELYS and that may be involved in NE-assembly and/or promote the formation of novel NPCs include the lamin B receptor (LBR; (Clever et al., 2012)) and the reticulon-like protein REEP4 (Golchoubian et al., 2022). In summary, a picture emerges, where ELYS serves as the crucial initiation factor for post-mitotic NPC-assembly. Remarkably, ELYS is not required for NPC-biogenesis during interphase (Hampoelz et al., 2019; Vollmer et al., 2015). In this study, we investigate a novel binding partner of ELYS, VAPB, in detail. Our results show that the MSP-domain of VAPB interacts with FFAT-like motifs of ELYS during mitosis in a phosphorylation-dependent manner. VAPB localizes to chromatin, in particular to the non-core region of the segregating chromosomes, together with ELYS, and depletion of VAPB resulted in prolonged mitosis and lagging chromosomes. Together, our findings suggest that VAPB serves as a factor that helps to recruit membranes and integral membrane proteins to chromatin during NE-assembly.

## Results

### The MSP-domain of VAPB mediates its interaction with FFAT motifs of ELYS

We recently identified the nucleoporin ELYS (AHCTF1) as a proximity partner of VAPB using RAPIDS (James et al., 2019). Furthermore, ELYS was identified as a potential interacting partner of VAPs in other proteomic studies (Huttlin et al., 2015; Murphy and Levine, 2016). We therefore decided to investigate the interaction of VAPB and ELYS in detail. First, we analyzed ELYS for potential FFAT motifs that would bind to the MSP-domain of VAPB. Three FFAT motifs were predicted in ELYS based on a position weight matrix strategy to identify short linear motifs (Di Mattia et al., 2020; Murphy and Levine, 2016; Slee and Levine, 2019), all of which lack the hydrophobic phenylalanine residues. FFAT-1 lies in the α-solenoid domain of ELYS, whereas FFAT-2 and FFAT-3 are found in its C-terminal region, which is predicted to be unstructured (Fig. 1A). FFAT-2, largely corresponds to a conventional FFAT motif with an aspartic residue in the fourth position of the core. FFAT-1 and FFAT-3, on the other hand, resemble phospho-FFAT motifs, as they contain a serine or threonine residue, respectively, at the fourth position. These residues could be phosphorylated to promote binding to VAPB (Di Mattia et al., 2020). In order to determine whether the three FFAT-motifs are indeed interaction sites for VAPB, we used a peptide binding approach (Di Mattia et al., 2020). Biotinylated peptides (Fig. 1B) were immobilized on beads and incubated with a total HeLa-cell lysate. For FFAT-1 and FFAT-3, phosphorylated and non-phosphorylated versions were used (pS_825_ and pT_1717_, respectively). A random peptide (Di Mattia et al., 2020) was used as negative control. As shown in Fig. 1C, strong binding of VAPB was detected for the FFAT-2 peptide. Much weaker binding was observed for the non-phosphorylated FFAT-1 peptide, whereas the phosphorylated version showed somewhat stronger binding. No binding of VAPB to the FFAT-3 peptides and the control peptide was observed. Next, we performed similar binding assays using immobilized peptides and GST-tagged versions of the MSP-domain of VAPB instead of cellular lysates. MSP-K87D/M89D (KD/MD) is an established mutant of VAPB that abolishes the interaction with FFAT-motifs (Kaiser et al., 2005). Importantly, the mutations do not affect folding and do not lead to protein aggregation (Kaiser et al., 2005). MSP-K43L, on the other hand, was reported as a mutant of VAPB that abrogates binding to phospho-FFAT motifs, but not to conventional FFAT-motifs (Di Mattia et al., 2020). As an additional control, we also used a peptide derived from ORP1 (also known as OSBPL1A), a protein with a strong FFAT motif (Murphy and Levine, 2016). Similar to the results shown in Fig. 1C, binding of the wild type MSP-domain of VAPB (GST-MSP) was stronger to the phosphorylated compared to the non-phosphorylated version of FFAT-1 (Fig. 1D). Strong binding was also observed for the FFAT-2 peptide, whereas FFAT-3 showed only background binding. As expected, the KD/MD mutation in the MSP-domain largely abrogated binding to FFAT-1 and FFAT-2 peptides and also to the ORP1-peptide. Surprisingly, the K43L-mutation clearly affected the interaction of GST-MSP with the strong FFAT motif of ORP1 as well. Accordingly, the K43L-mutation also affected binding of GST-MSP to the FFAT-1 and FFAT-2 peptides. Importantly, no binding of purified GST to any of the immobilized peptides was detected. Together, these data show that VAPB interacts with ELYS-peptides via its MSP-domain and that the interaction is mediated by different types of FFAT-motifs, which may in part be regulated by phosphorylation.

**Figure 1:**
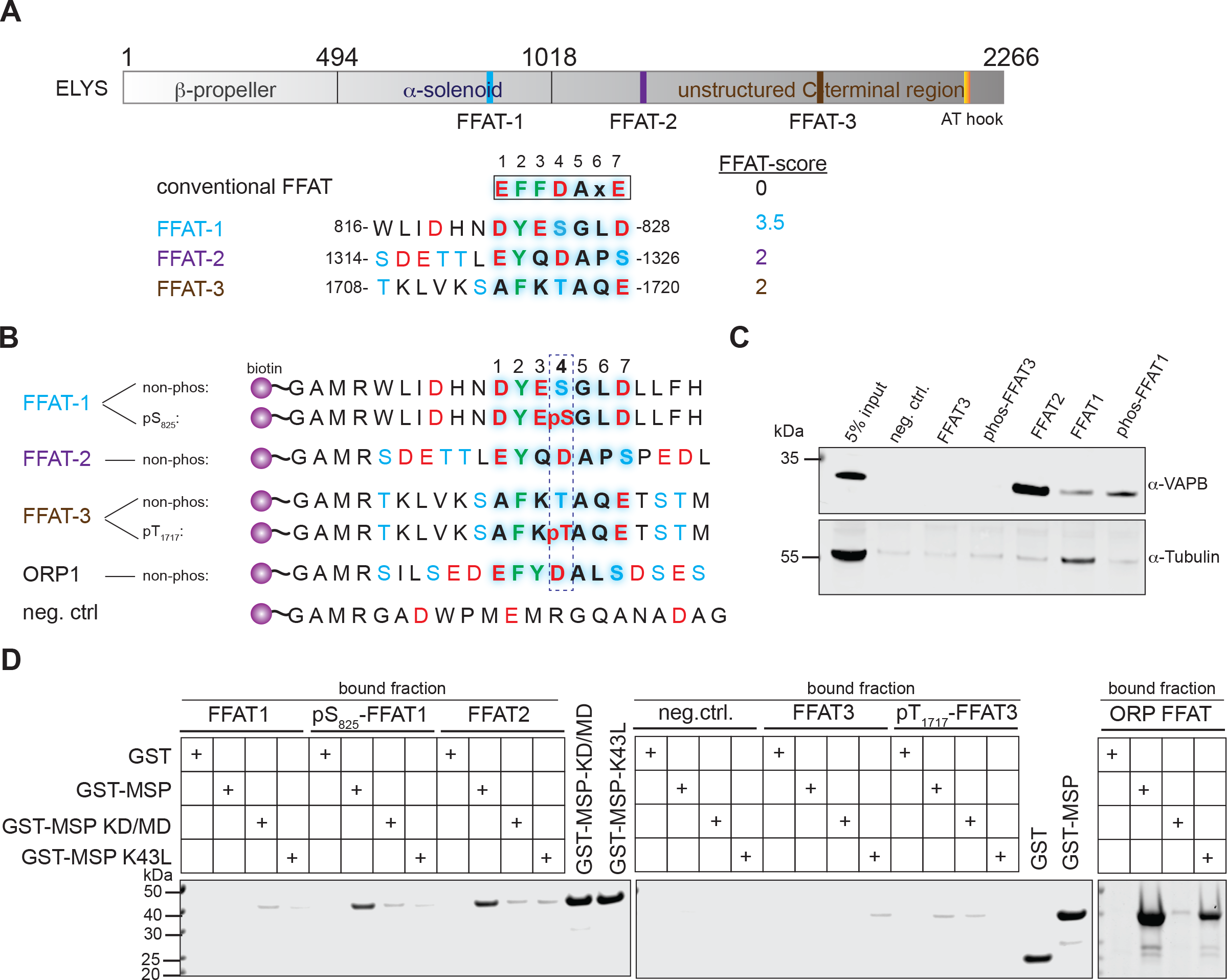
VAPB-ELYS interactions are mediated by FFAT- and phospho-FFAT motifs. (A) Schematic representation of the domain structure of ELYS. The predicted FFAT motifs are shown in blue, purple and brown with the respective sequences and FFAT scores (the lower the score, the more similar it is to the conventional FFAT motif) below (with six acidic tract residues and seven core residues). The AT-hook is shown in orange. (B) Sequences of ELYS-, OPR1- and negative control peptides used for pull down assays. Peptides contain biotin at the N-terminus, a linker sequence (GAMR) and the FFAT sequences of ELYS (FFAT-1 and FFAT-3, either with or without phosphorylated serine-(pS) or threonine (pT) residue at position 4 of the core motif, and non-phosphorylated FFAT-2). The ORP1-peptide (amino acid residues 469-485) was used as a positive and a random peptide as a negative control. (C) Peptides as in B were immobilized on beads and incubated with total HeLa-cell lysate. Interacting proteins were analyzed by SDS-PAGE, followed by Western blotting using antibodies against VAPB or tubulin. (D) Peptides as in B were immobilized on beads and incubated with purified proteins (GST, the wild type MSP-domain of VAPB (GST-MSP), the KD/MD mutant (GST-MSP-KD/MD) or the K43L mutant of the MSP-domain (GST-MSP-K43L). Interacting proteins were analyzed by SDS-PAGE and Coomassie-staining. 10% of the input of the recombinant proteins were loaded as indicated.

### Phosphorylation of ELYS regulates VAPB binding

Our findings described above show that the MSP-domain of VAPB is involved in binding to isolated ELYS-peptides. We next investigated binding of the endogenous full-length nucleoporin to overexpressed or endogenous VAPB. Many nucleoporins are extensively phosphorylated at the onset or during mitosis (Blethrow et al., 2008; Glavy et al., 2007; Macaulay et al., 1995). Although multiple phosphorylation sites have been identified in ELYS (Hornbeck et al., 2015), potential consequences of ELYS-phosphorylation have not been investigated. First, we generated a cell line expressing GFP-VAPB in a tetracycline-inducible manner. Cells were enriched in the G1/S-phase of the cell cycle or in mitosis, or left unsynchronized (asyn) and subjected to a co-immunoprecipitation protocol using nanobodies (GFP-Selector). As shown in Fig. 2A, a large proportion of GFP-VAPB was precipitated under our experimental conditions. ELYS could not be detected as a coprecipitating protein when a lysate from asynchronous cells was used. Lysates from G1/S-cells and from mitotic cells, by contrast, yielded a small amount of co-precipitating ELYS. ELYS from the mitotic lysate migrated as two distinct bands on the SDS-gel (Fig. 2A, input), suggesting a post-translational modification of the protein. ELYS was not detected when the cells had not been induced with tetracycline to express GFP-VAPB. Next, we performed an immunoprecipitation experiment using antibodies against endogenous VAPB, now using lysates from asynchronous and mitotic cells. Similar to the results in Fig. 2A, the signal of co-precipitating ELYS was more pronounced for the mitotic lysate (Fig. 2B). When purified IgG was used as a control, only background levels of ELYS were detected. In a next step, we analyzed binding of endogenous ELYS to immobilized versions of the MSP-domain of VAPB, purified from bacteria as GST-fusion proteins. As shown in Fig. 2C, strong binding of ELYS to wild-type GST-VAPB-MSP was observed when a mitotic lysate was used. Binding was much weaker with a lysate from asynchronous cells and no binding to immobilized GST was detected. As expected for a typical MSP-based interaction, binding of ELYS was abolished when the KD/MD-version of GST-VAPB-MSP was used as an immobilized protein. Finally, we directly addressed the question of a possibly phosphorylation-dependent interaction of VAPB and ELYS. We first treated our lysates with λ-phosphatase to dephosphorylate cellular proteins. Indeed, this treatment resulted in an increased electrophoretic mobility of ELYS in the mitotic lysate (Fig. 2D; input). Phosphorylation of ELYS could also be confirmed using phos-tag gels (data not shown). Lysates that had been treated with or without λ-phosphatase were then incubated with immobilized GST-VAPB-MSP. Both, the weak interaction seen with ELYS from a lysate from asynchronous cells and the much stronger interaction seen with ELYS from a mitotic lysate was abolished upon λ-phosphatase treatment (Fig. 2D). Together, these results clearly show that VAPB and ELYS interact with each other most prominently during mitosis and that this interaction can be regulated by phosphorylation.

**Figure 2:**
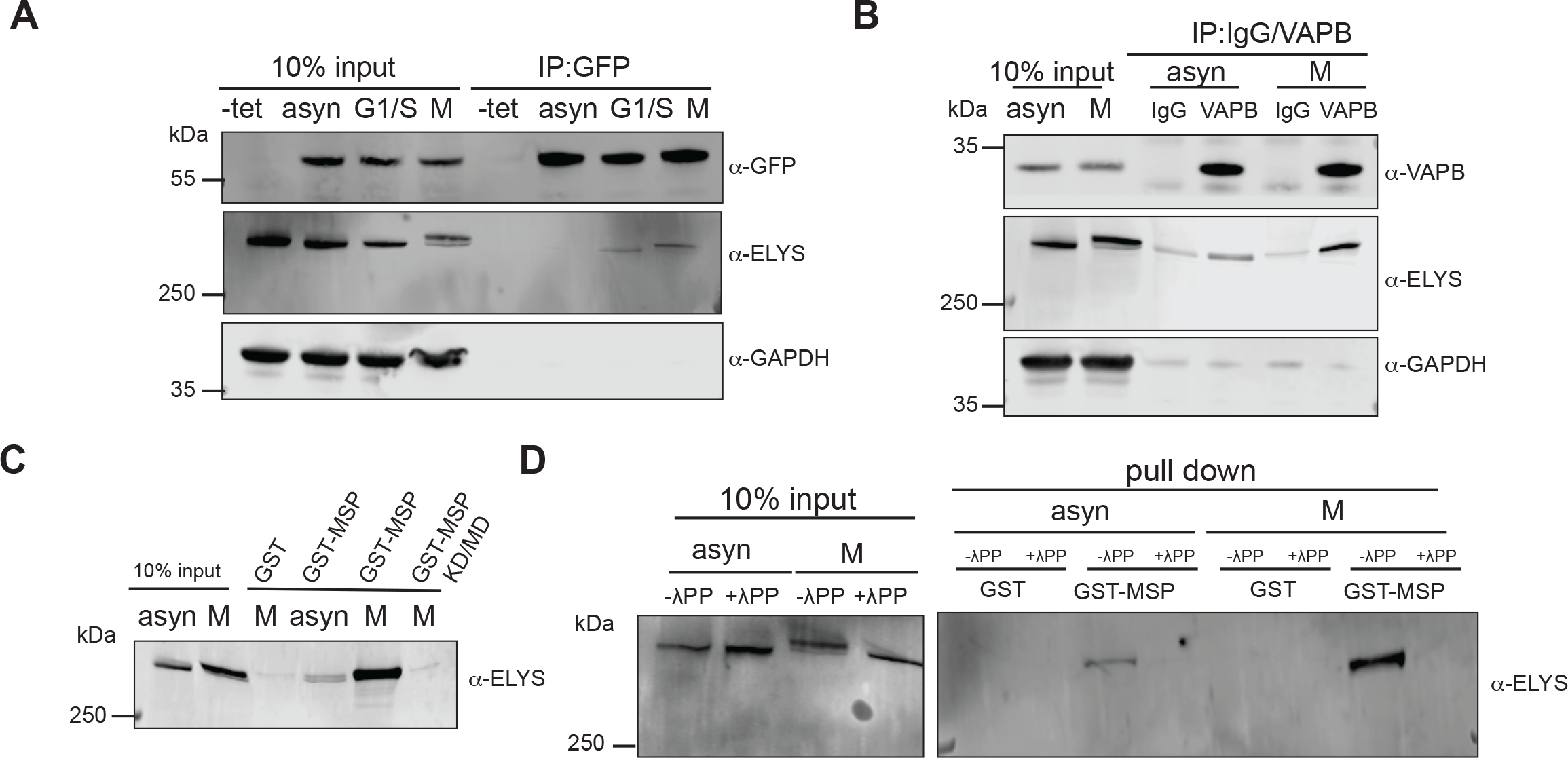
Binding of VAPB to ELYS is regulated by phosphorylation. (A) HeLa Flp-In Trex cells with stable expression of GFP-VAPB were treated with or without (-tet) tetracycline to induce expression. Cells were then synchronized either in the G1/S phase by a double thymidine block or in mitosis (M) by treatment with DME. Total cell lysates obtained from asynchronous (asyn) or synchronized (G1/S and M) cells were subjected to co-immunoprecipitation (IP) using the GFP-Selector. (B) HeLa cells were treated as in A to obtain asynchronous or mitotic lysates, which were subjected to co-IP using antibodies against VAPB or purified IgG as a control. A and B, precipitated proteins were analyzed by SDS-PAGE, followed by Western blotting using antibodies against GFP, ELYS, VAPB and GAPDH as indicated. Note the slower migrating form of ELYS in mitotic lysates. (C) Purified proteins (GST, the wild type MSP-domain of VAPB (GST-MSP) and the KD/MD mutant of MSP (GST-MSP-KD/MD) were immobilized on GST-Selector beads and incubated with lysates from asyncronous (asyn) or mitotic (M) HeLa cells. (D) Lysates from asyncronous (asyn) or mitotic (M) HeLa cells were treated with (+) or without (-) /\-phosphatase (/\PP) and incubated with purified proteins (GST or GST-MSP) bound to GST Selector agarose beads. (D) and (E) Interacting proteins were analyzed by SDS-PAGE, followed by Western blotting using antibodies against ELYS.

### FFAT-2 of ELYS is regulated by phosphorylation

In light of the strong dependency of VAPB-binding to ELYS on phosphorylation, we decided to characterize the phosphorylation state and potential phosphorylation sites of ELYS in more detail. Cell lysates from asynchronous or mitotic cells were incubated with GST-VAPB-MSP immobilized on beads. As before (Fig. 2B), the mitotic lysate yielded more protein in the high-molecular weight range compared to the lysate from asynchronous cells (Fig. 3A). Proteins were then subjected to mass spectrometry for the identification of phosphorylated peptides (Fig. 3A). Fig. 3B shows the phosphorylation sites on ELYS identified by our approach, and also sites that had previously been reported by others (Hornbeck et al., 2015; Sharma et al., 2014; Ullah et al., 2016). Despite a total of 156 identified phosphorylation sites, our candidate phospho-FFAT-motif (FFAT-1) was not identified as a hit. FFAT-2, however, is close to a phospho-cluster and also contains two serine residues that can be phosphorylated, serine 1314 and serine 1326. We therefore decided to look in more detail into potential effects on VAPB-binding, depending on the phosphorylation status of these sites. Following the approach as described in Fig. 1, several biotinylated versions of the FFAT-2 peptide were synthesized: the non-phosphorylated version (as in Fig. 1), two phosphorylated versions with phospho-serine residues at positions 1314 and 1326, respectively, a double-phospho-version with both residues phosphorylated and one version where both serines are changed to alanines (Fig. 3C). Again, peptides were immobilized on beads, incubated with a total-cell lysate and interacting proteins were analyzed. Figs. 3D and E clearly show that phosphorylation of serine 1314, which is upstream of the core region of the FFAT motif, strongly promotes binding of VAPB to the peptide. Phosphorylation of the core residue serine 1326, by contrast, slightly reduced binding of VAPB, in particular when serine 1314 was phosphorylated. Likewise, the alanine-containing peptides allowed only very little binding of endogenous VAPB. In a next step, we analyzed binding of purified proteins to the immobilized peptides, using GST and the wild-type- and the KD/MD-version of the MSP-domain of VAPB as potential interaction partners. Similar to the results described above, phosphorylation of serine 1314, but not serine 1326 of the FFAT-2 motif enhanced binding of the wild-type MSP-fusion protein to the peptide. Only background binding was observed when the mutant version of the VAPB-MSP domain (KD/MD) was used, demonstrating the specificity of the interaction. Together, these results suggest that phosphorylation of FFAT-2, as it may occur in particular during mitosis, can regulate binding of VAPB to ELYS. To what extent phosphorylation of the many other phospho-sites in ELYS contribute to the phosphorylation-dependent interaction of the two proteins (Fig. 2D) remains to be investigated.

**Figure 3:**
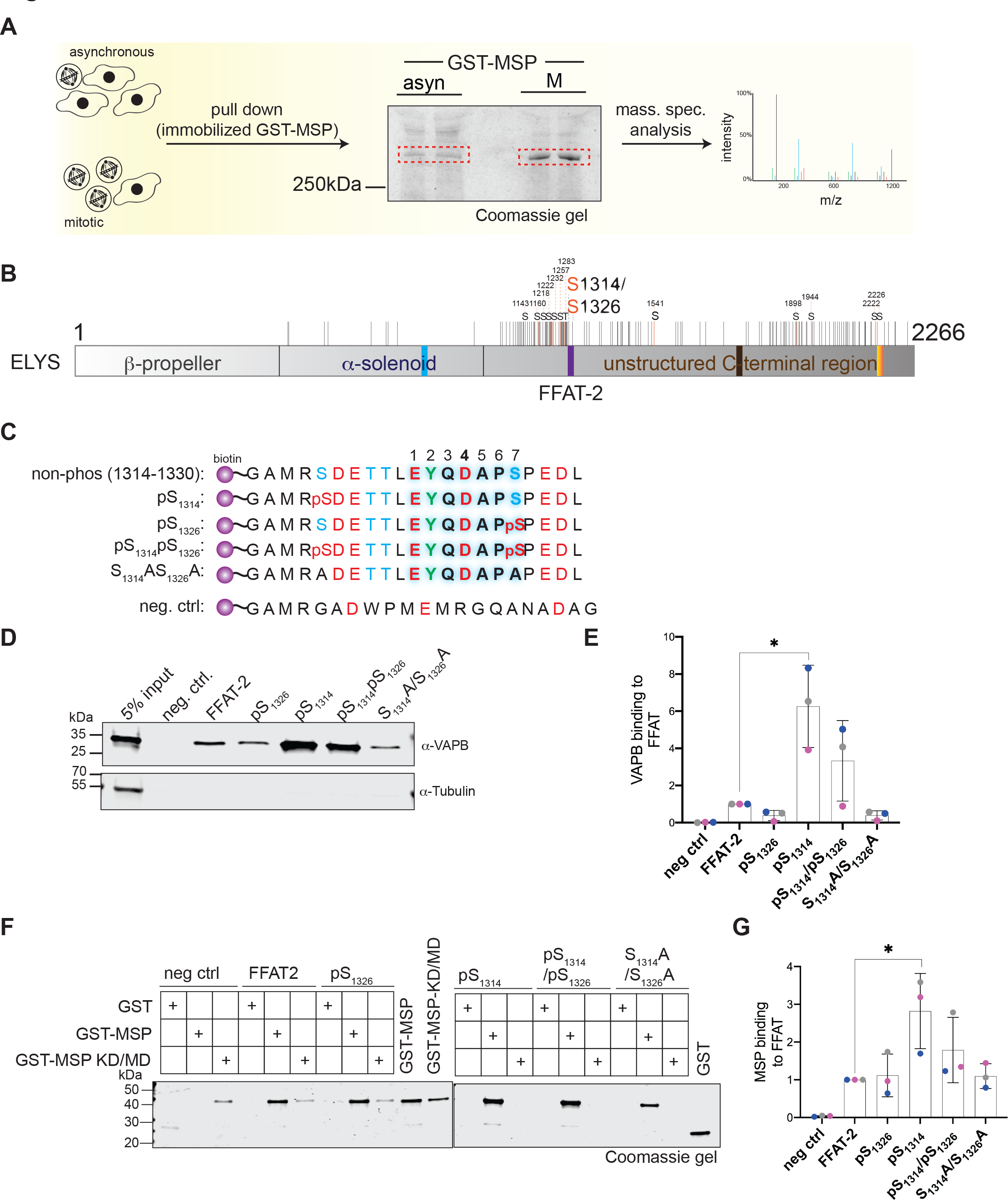
Phosphorylation of serine 1314 in the acidic tract of FFAT-2 enhances the interaction of ELYS with VAPB. (A) Lysates from asynchronous and mitotic HeLa cells were subjected to pull down reactions using the MSP-domain of VAPB bound to GST Selector (compare Fig. 2C). Bound proteins were detected using Coomassie staining. The bands above 250 kDa (represented by rectangular region) were excised and analyzed by mass spectrometry to identify phosphorylation sites. (B) Phosphorylation sites identified by mass spectrometry (orange lines) as well as sites that had been identified before (Hornbeck et al., 2015; Sharma et al., 2014; Ullah et al., 2016); black lines) are indicated on the ELYS protein. Two phospho-serine sites (S1314 and S1326) are within the FFAT-2-motif. (C) Sequences of the ELYS-peptides used for pull down assays. Peptides contain biotin at the N-terminus, a linker sequence and the FFAT-2 motif either with or without phosphorylated serine-residues at position 1314 and/or 1326 or alanine residues at these positions. A random peptide served as negative control. (D) Peptides as in C were immobilized and incubated with total HeLa cell lysates. Bound proteins were analyzed by SDS-PAGE, followed by Western blotting using antibodies against VAPB and tubulin. (E) Quantification of three independent experiments as in (D). VAPB-binding to FFAT-peptides was normalized to non-phosphorylated FFAT-2. (F) Peptides as in C were immobilized and incubated with GST or GST-tagged wild type (GST-MSP) or the mutant version (GST-MSP KD/MD) of the MSP-domain. Bound proteins were analyzed by SDS-PAGE and Coomassie staining. The input fractions are loaded as indicated. (G) Quantification of three independent experiments as in (F). GST-MSP-binding to FFAT-peptides was normalized to non-phosphorylated FFAT-2.

### VAPB co-localizes with ELYS in anaphase

We have previously shown that ELYS and VAPB are found in close proximity at the INM of interphase cells (James et al., 2019). ELYS is known to localize to kinetochores in metaphase (Rasala et al., 2006) and then, later in mitosis, to the decondensing chromosomes, where it functions as an interaction platform for the formation of novel NPCs (Franz et al., 2007; Hampoelz et al., 2019). The localization of VAPB in mitosis has, to the best of our knowledge, not been analyzed before. We first used polyclonal antibodies against endogenous VAPB to detect the protein during different stages of mitosis. In interphase and in early prophase, VAPB was mainly found at the level of the ER and also at the NE (Fig. 4A), as shown before. In metaphase, VAPB was largely distributed over the cells, but was clearly absent from the chromatin region, where the mitotic spindle is thought to create a zone that excludes organelles and membrane structures from this area of the cell (Lu et al., 2009; Smyth et al., 2012). In anaphase and telophase, the protein was then detected adjacent to the chromatin, suggesting that it is present at the newly forming NE. In these stages of mitosis, the spindle microtubules are in contact with the “core region” of the segregating chromosomes, which also has a characteristic protein composition (Fig. 4B; for review see (Liu and Pellman, 2020)). ELYS and a number of other proteins, by contrast, are known to preferentially localize to the “non-core region” of the chromosomes. For a detailed analysis of the localization of VAPB at mitotic chromosomes, we made use of inducible cell lines expressing HA- or GFP-tagged version of VAPB. As shown in Fig. 4C, LBR was found in non-core regions of segregating chromosomes during anaphase, whereas emerin was detected mainly in core regions, as described previously (Haraguchi et al., 2000; Lee et al., 2001). For GFP-VAPB, we found a clear colocalization with ELYS at the non-core region. In a next step, we further characterized a possible interaction of ELYS and VAPB during mitosis using proximity ligation assays (PLAs (Soderberg et al., 2006)). As shown in Fig. 4D, we could detect the strongest PLA-signals for the ELYS-HA-VAPB pair in anaphase cells, whereas other mitotic stages and interphase cells yielded much lower signals. In PLAs, tagged versions of a protein of interest offer the possibility to compare wild-type proteins with proteins containing pertinent mutations at key residues. As shown above (Figs. 1D, 2C and 3F), the KD/MD-mutation in the MSP-domain of VAPB largely abolishes interaction with ELYS. We therefore compared PLA-signals around the chromosome masses in anaphase cells expressing either wild-type full-length HA-VAPB or the mutant HA-VAPB-KD/MD. Clearly, the mutant protein yielded less PLA-interactions compared to wild-type VAPB, suggesting a direct interaction of VAPB and ELYS at the segregating chromosomes in anaphase (Fig. 4E, F). The rather small difference observed between the wild-type and the mutant protein observed in this experiment probably results from the presence of endogenous VAPB in the stable cell lines, which could form dimers with the exogeneous HA-tagged versions. Hence, also HA-VAPB-KD/MD can be detected in proximity to endogenous ELYS to some extent.

**Figure 4:**
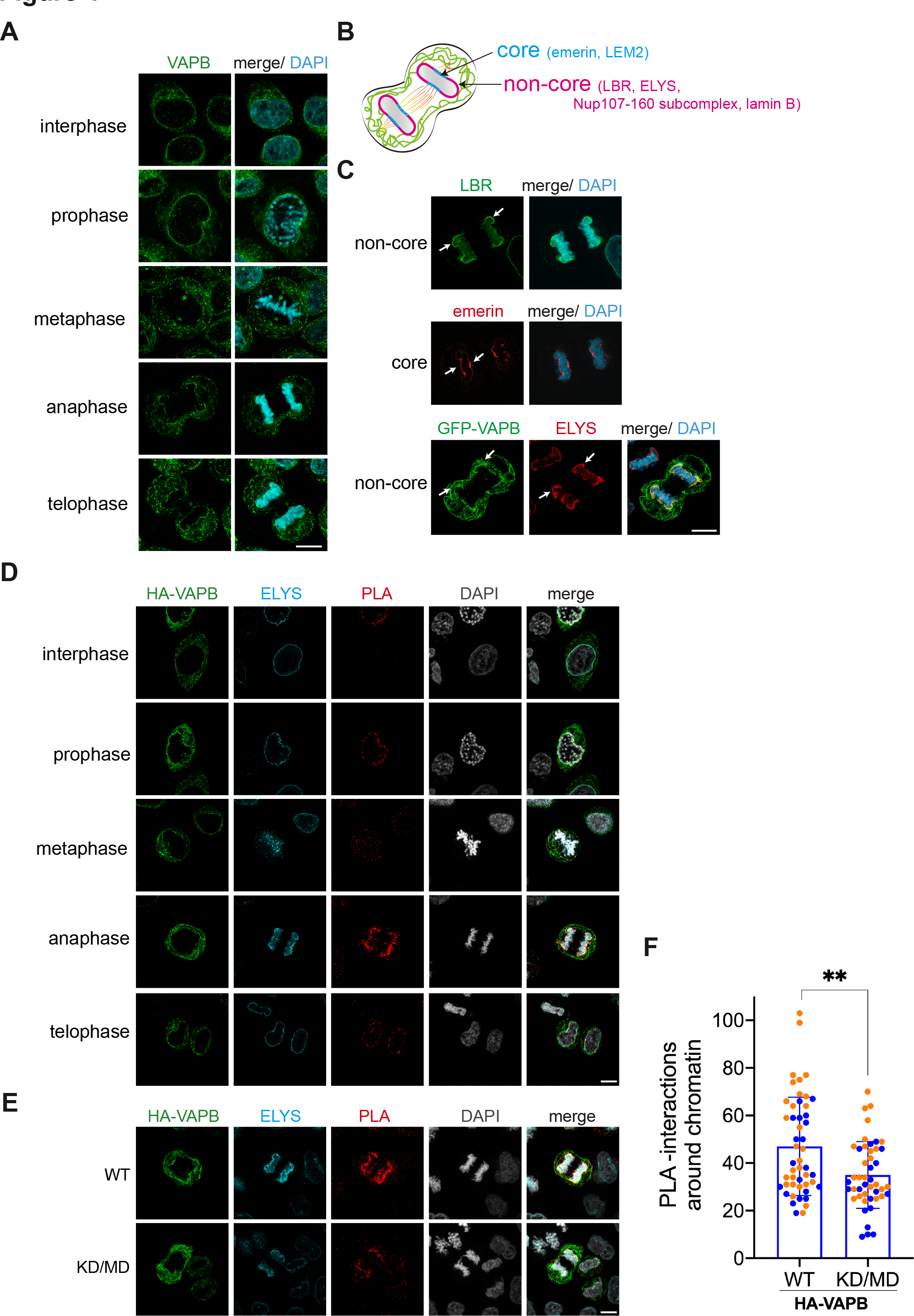
VAPB localizes to the non-core regions on mitotic chromosomes. (A) Intracellular localization of VAPB in interphase, prophase, metaphase, anaphase and telophase. HeLa P4 cells were synchronized by double thymidine block and released for 9 h before fixing and staining with antibodies against VAPB. (B) Schematic representation of anaphase cell with core (blue) and non-core (magenta) nuclear envelope (NE) subdomains. Green lines indicate ER-membranes. (C) HeLa Flp-In Trex cells stably expressing GFP-VAPB were synchronized and released as in A and subjected to indirect immunofluorescence staining using antibodies against LBR and emerin. Arrows indicate the core and non-core NE subdomains. (D) HeLa Flp-In Trex cells stably expressing HA-VAPB were synchronized and released from thymidine block as described in A and subjected to PLAs using antibodies against HA and ELYS. HA-VAPB and ELYS were detected by indirect immunofluorescence. (E) After synchronization and release, Flp-In Trex cells stably expressing either wild type (WT) or the KD/MD mutant of HA-VAPB were subjected to PLAs using antibodies against HA and ELYS as described in D. Combined z-stacks of cells in anaphase are shown. (A, C-E) DNA was stained with DAPI and cells were analyzed by confocal microscopy. Scale bars, 10 µm. (F) Quantification of PLA interactions detected around chromatin, analysing a total of 24 anaphase cells (i.e. 2n chromatin regions). **, p<0.01

### Mitotic defects in VAPB-depleted cells

The specific localization of VAPB at mitotic chromosomes suggested that the protein might also have a function during this stage of the cell cycle. We therefore addressed this issue and investigated key mitotic parameters in control cells and in VAPB-depleted cells. To monitor the progression through mitosis, cells that had been treated with siRNAs against VAPB or with non-targeting siRNAs (Fig. 5A) were first subjected to a synchronization protocol, involving a double-thymidine block and the reversible kinesin inhibitor dimethylenastron (DME) (Muller et al., 2007), which induces a mitotic arrest in prometaphase (Ertych et al., 2014) (Fig. 5B). After release from the DME-arrest, cells were harvested every 15 minutes for a total of 2 h and analyzed by flow cytometry. As shown in Figs. 5C and D, VAPB-depleted cells displayed a slower progression through mitosis compared to the control cells, with a clear delay right after the release. For a more detailed analysis of possible perturbations during mitosis in VAPB-depleted cells, we resorted to a live-cell imaging approach, using cells expressing an mCherry-tagged version of histone H2B. Cells that had been treated with siRNAs against VAPB or with non-targeting siRNAs were subjected to a double-thymidine block, released and imaged to capture mitotic cells. Strikingly, about 90% of the VAPB-depleted cells exhibited a chromosome segregation defect with lagging chromosomes in anaphase, compared to only 10% of the control cells (Figs. 5E and F and Videos 1 and 2). A detailed inspection of the lengths of individual mitotic phases further revealed that the depleted cells are clearly slowed down with respect to the transition from meta-to anaphase (Fig. 5G). Other mitotic phases were not affected by the siRNAs against VAPB. In summary, our results suggest a scenario, where phosphorylation-dependent binding of VAPB-containing membrane fragments to chromatin-associated ELYS contributes to a timely formation of a novel NE during mitosis.

**Figure 5:**
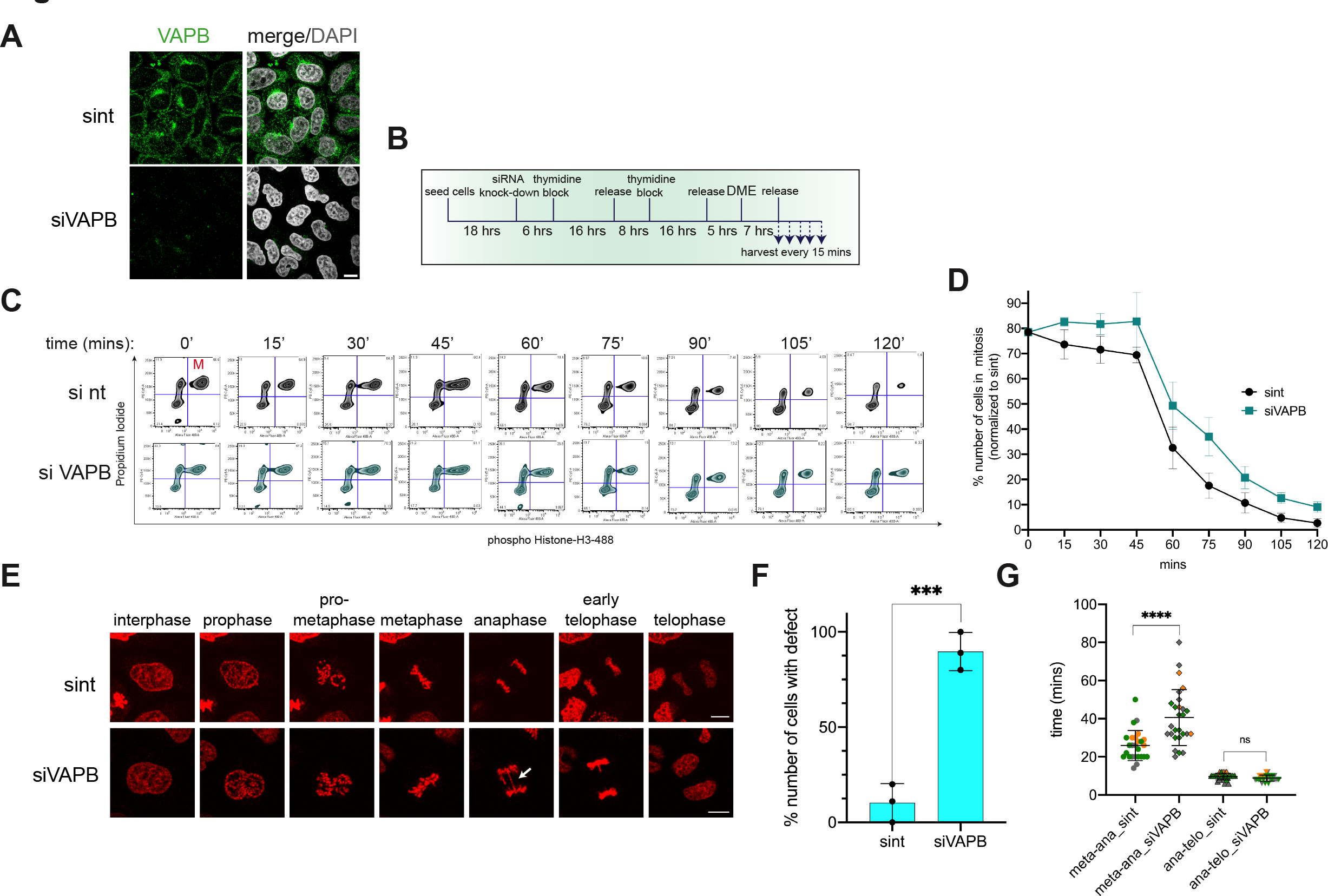
Depletion of VAPB delays mitosis and leads to chromosome segregation defects. (A) HeLa P4 cells were treated with siRNAs against VAPB (siVAPB) or non-targeting siRNAs (sint) and analyzed by indirected immunofluorescence, detecting endogenous VAPB, and confocal microscopy. Scale bar, 10 µm. (B) Schematic representation of synchronization- and release steps for HeLa P4 cells after knocking down VAPB using siRNA. Cells were transfected using either non-targeting or siRNA against VAPB. (C) HeLa P4 cells were transfected with siRNAs against VAPB (siVAPB) or non-targeting siRNAs (sint) and synchronized and further treated as depicted in A. Cells were then analyzed by flow cytometry using propidium iodide and antibodies against phospho-Histone H3. Gated mitotic cells are observed in the upper right quadrant. (D) Quantification of three experiments as in B, depicting the percentage of mitotic cells as determined by flow cytometry. Error bars indicate the variation from the mean of three independent experiments. (E) Live cell imaging of HeLa Flp-In Trex cells stably expressing mCherry-H2B after knock down of VAPB. Cells transfected with either non-targeting siRNAs (sint) or siVAPB were synchronized by a double thymidine block. After releasing the block for 6 h, the cells were imaged every 2 min for a total of 2 h after the onset of mitosis. Examples from different phases of mitosis are shown. The arrows indicate lagging chromosomes. Scale bar, 10 µm. See also Videos 1 and 2. (F) Cells were treated as in D and the percentage of cells showing chromosome segregation defects was analyzed. Error bars indicate the standard deviation from the mean of three independent experiments. ***, p<0.001. (G) Cells were treated as in D and the progression from metaphase to anaphase and from anaphase to telophase was measured using live cell imaging. Three independent experiments were performed, analyzing a total of 26 cells for si nt and 27 cells for siVAPB. ****, p<0.0001; ns, not significant.

## Discussion

VAPB has been extensively studied as an ER-resident protein that serves as a binding partner for other membrane proteins, supporting the formation and/or maintenance of cellular contact sites between the ER and other organelles (James and Kehlenbach, 2021). Here we describe the unexpected interaction of VAPB, which is also found at the INM, with ELYS, a nucleoporin that does not contain a transmembrane domain. Instead, ELYS directly interacts with chromatin during mitosis via its AT-hook and recruits nucleoporin-subcomplexes to initiate the formation of novel NPCs. Concurrent with the separation of chromosomes during anaphase, membranes derived from the mitotic ER assemble around the chromatin masses to ultimately form a novel NE ((Anderson and Hetzer, 2007; Anderson et al., 2009), for review see also (Guttinger et al., 2009)). This process has to be tightly coordinated with the biogenesis of the NPC and we suggest that VAPB has a function here, as it interacts with ELYS, particularly in anaphase cells.

### Phosphorylation-dependent binding of VAPB to ELYS

Many nucleoporins and also nuclear lamins (Gerace and Blobel, 1980; Kutay et al., 2021) are known to be phosphorylated at the onset of mitosis. Phosphorylation of Nup98, for example, is a rate-limiting step in the disassembly of the NPC (Laurell et al., 2011). Our results clearly show, that ELYS is also heavily phosphorylated during mitosis, indicated by its distinct shift in mobility in SDS-PAGE in mitotic lysates compared to lysates from cycling cells (Fig. 2). In light of the very large total number of phosphorylation sites in ELYS (>150), it will be very difficult to determine the role of individual sites on mitotic progression. We therefore focused on sites that are part of the predicted FFAT motifs of ELYS to investigate in detail possible consequences on VAPB-ELYS interactions. What is the basis for these interactions? Our initial biotinylation-approach (James et al., 2019) revealed ELYS as a *proximity*-partner of VAPB and did not discriminate between direct and indirect interactions. Using purified protein fragments of VAPB and immobilized ELYS-peptides, we now show that the proteins can indeed interact with each other, without the need for additional bridging factors (Figs. 1 and 3). Furthermore, our co-immunoprecipitation experiments (Fig. 2) combined with peptide-binding assays clearly show that the interaction is regulated by phosphorylation. Of particular interest here is a sequence (FFAT-2) in ELYS that was identified as an FFAT-like motif, as phosphorylation of a serine residue (S1314) in the acidic tract strongly promoted binding of the MSP-domain of VAPB. Another serine residue in the core region (S1326) had also been identified as a phospho-site in ELYS. In our assays, phosphorylation of this amino acid residue resulted in reduced binding of endogenous VAPB (Figs. 3D, E) or the purified MSP-domain of VAPB (Figs. 3G, F). Examples for such a regulation by phosphorylation within FFAT motifs have been described before (Kors et al., 2022). Overall, phosphorylation clearly promoted binding of VAPB to ELYS, as treatment of lysates with /\-phosphatase largely abrogated the interaction. FFAT-2 is found in a region of ELYS without a predicted structure, as it is typically the case for FFAT motifs (Mikitova and Levine, 2012). Remarkably, this region is highly conserved between species, suggesting that it is important for protein functions (data not shown). Another motif, FFAT-1, which is found in a structured region of ELYS, was initially identified with an FFAT-score of 3.5, indicating a comparatively low level of similarity to the conventional FFAT-sequence. Nevertheless, we detected binding of endogenous VAPB and of purified MSP-domains to the corresponding peptide (Fig. 1), which was further enhanced upon phosphorylation of a serine residue (S825). It remains to be investigated, how phosphorylation of two or more residues affects the interaction with VAPB in the context of full-length ELYS. Interestingly, ELYS also interacts with protein phosphatase 1 (PP1; (Hattersley et al., 2016)), which may affect the phosphorylation status of adjacent proteins like LBR (Mimura et al., 2016). The proposed docking site for PP1 on ELYS is very close to its FFAT-2 motif, but it is unknown if PP1-activity is required for de-phosphorylation of ELYS itself.

### Mitotic roles of VAPB

Binding of ELYS to chromatin is an early event in post-mitotic NPC-biogenesis (Franz et al., 2007; Rasala et al., 2008) that is followed by the recruitment of components of the Y-complex (Franz et al., 2007; Gillespie et al., 2007). In the complete absence of ELYS, nuclear envelopes can be formed but lack NPCs entirely (Franz et al., 2007). It was also reported that further oligomerization of original nucleoporin subcomplexes requires the recruitment of membrane components to the site of NPC-assembly (Rotem et al., 2009) and that nuclear envelope formation always precedes NPC-assembly (Lu et al., 2011), although alternative scenarios have been suggested (for review see (Liu and Pellman, 2020)). Whatever the exact sequence of events, integral membrane proteins are expected to play important roles in these steps. Indeed, several proteins of the INM can directly interact with chromatin, for example LBR (Ye and Worman, 1994)) and Lap2β (Foisner and Gerace, 1993). Furthermore, INM-proteins in general are enriched in basic domains that may facilitate membrane interactions with DNA (Ulbert et al., 2006). Our data now suggest that VAPB can also function in directing membranes to chromatin, with ELYS as a bridging factor. VAPB localized largely to the non-core region of anaphase chromosomes, together with ELYS. Depletion of VAPB resulted in prolonged mitosis, with particularly lengthened transition times from metaphase to anaphase (Fig. 5), i.e. at stages in mitosis where the close proximity/interaction of ELYS and VAPB was most pronounced (Figs. 4D-F). Similar effects have been observed upon knock down of several INM proteins like LBR, Lap2β and MAN1. However, reduction in any one of these proteins did not block NE formation and is thus indicative of a redundant system (Anderson et al., 2009). A delay at individual stages, however, can lead to chromosome segregation defects, as seen for VAPB (Figs. 5E, F), with lagging chromosomes or chromosome bridges. Ultimately, such defects could also lead to the generation of micronuclei and genome instability, although this was not observed in our studies (see Videos 1 and 2). Hence, it appears that the cells are able to cope with the lack of VAPB, allowing more time for a faithful segregation of chromosomes and thus avoiding deleterious effects. Similar observations were made in cells overexpressing Nup107 or lamin B, which enforced a chromosome separation checkpoint to prevent NE reformation on incompletely separated chromosomes (Afonso et al., 2014). Clearly, a function of VAPB in mitosis is not essential, as VAPB-knockout cells (Dorsch et al., 2021) as well as knockout mice (Silbernagel et al., 2018) are viable. One cellular component that could function here as a rescue factor is VAPA, a protein very similar in sequence and localization to VAPB (Lev et al., 2008). In our experiments, knockdown of VAPA alone did not lead to a delay in mitosis (data not shown). Hence, it seems likely that completely independent mechanisms apply as well, relying on different proteins that mediate and coordinate the interaction of membranes with chromatin. The large number of INM proteins with the ability to function in membrane-recruitment may provide a safety mechanism in post-mitotic NE-formation, where loss of a single protein may affect the efficiency of the process but is overall tolerated.

## Materials and methods

### Plasmids

For pcDNA5/FRT/TO-GFP, the GFP sequence was amplified by PCR using primers G2594 and G2595 and pEGFP-C3 (addgene #2489) as template (oligonucleotides are listed in Table 1). The PCR product was cloned into pcDNA5-FRT-TO vector (Thermo Fischer Scientific) via AflII and HindIII. The VAPB coding sequence was amplified using primers G1390 and G1386 and pCAN-myc-VAPB (Fasana et al., 2010) as a template and inserted into pcDNA5/FRT/TO-GFP via KpnI and BamHI. To obtain pcDNA5/FRT/TO-HA-VAPB, the HA-tag was inserted into pcDNA5/FRT/TO by oligo annealing using oligos G2599 and G2600. The VAPB coding sequence was amplified using primers G2601 and G2602 and pCAN-myc-VAPB (Fasana et al., 2010) as a template and inserted into pcDNA5/FRT/TO-HA via EcoRV and NotI. Site-direct mutagenesis of pcDNA5/FRT/TO-HA-VAPB was performed using primers G2403, G2404, G2405, G2406 to generate pcDNA5/FRT/TO-HA-VAPB-K87D/M89D.

**Table 1:**
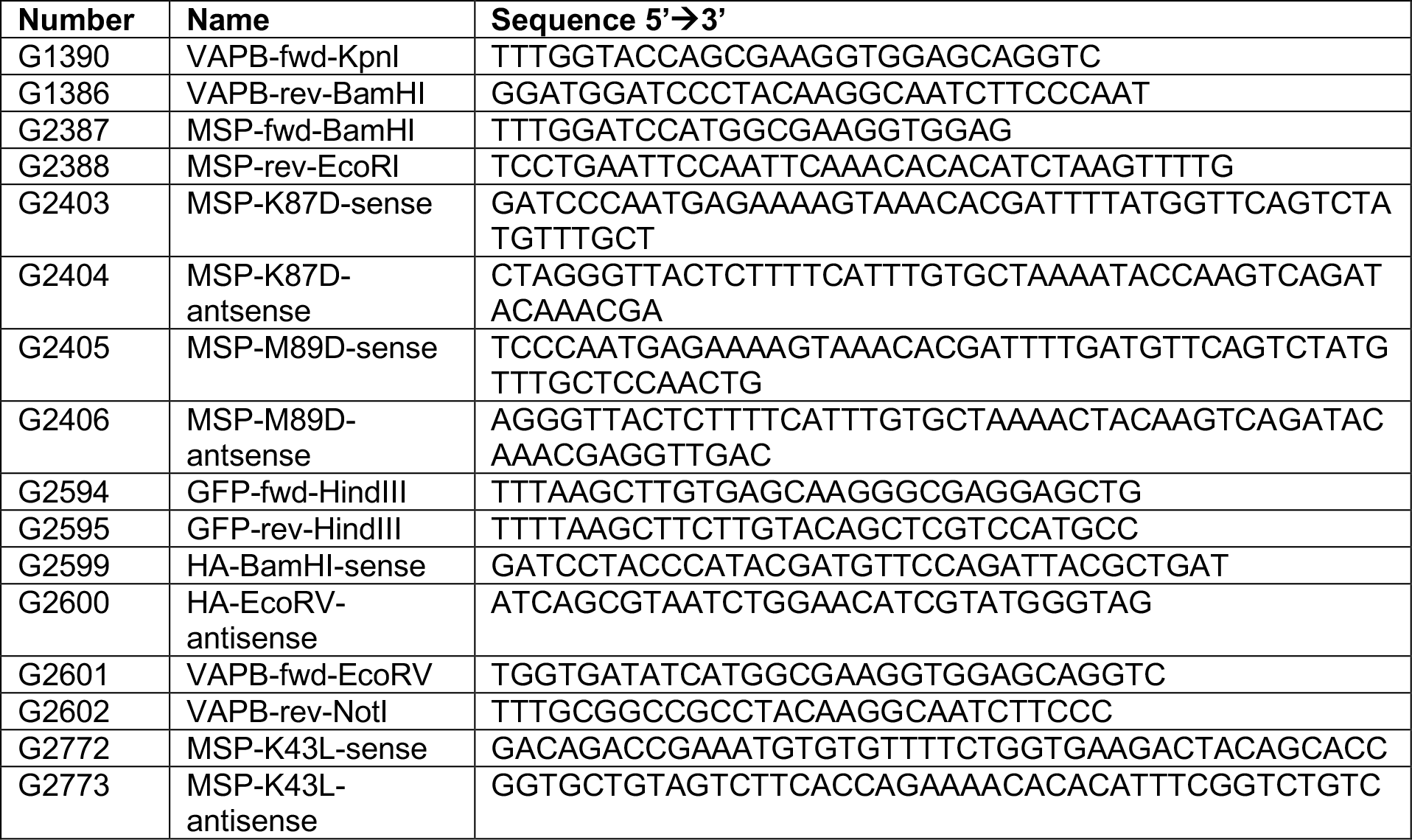
Oligonucleotides used for cloning.

For pGEX-6P-1-MSP-VAPB, the MSP domain of VAPB was amplified by PCR using primers G2387 and G2388 and pCAN-myc-VAPB (Fasana et al., 2010) as a template. The PCR product was cloned into pGEX-6P-1 through EcoRI and BamHI. To generate the MSP-KD/MD mutant, site-directed mutagenesis was performed as described above using primers G2403, G2404, G2405, G2406. To generate the MSP-K43L mutant, site-directed mutagenesis was performed using primers G2772 and G2773. All plasmids obtained were confirmed by sequencing (Eurofins Genomics).

### Cell culture, transfection and drug treatment

Parental HeLa Flp-In T-REx (Hafner et al., 2014) and H2B-cherry HeLa Flp-In T-REx cells used to generate stable tetracycline-inducible cell lines were a gift from T. Mayer (University of Konstanz, Germany). HeLa P4 cells (Charneau et al., 1994) were obtained from the NIH AIDS Reagent Program. Cells were grown in DMEM supplemented with 10% (v/v) normal or tetracycline-free FBS (Thermo Fischer Scientific), 100 U ml^-1^ penicillin, 100 µg ml^-1^ streptomycin and 2 mM l-glutamine (Thermo Fischer Scientific) at 37 °C in the presence of 5% CO_2_ and tested for contamination by mycoplasma on a regular basis.

siRNA-mediated knockdown of VAPB was carried out using Lipofectamine RNAiMAX (Thermo Fisher Scientific), following instructions of the manufacturer. VAPB siRNAs (GCUCUUGGCUCUGGUGGUUUU and AAAACCACCAGAGCCAAGAGC; Sigma) and ON-Target-plus nontargeting siRNA (D-001810-01-50, Dharmacon, Lafayette, CO) were used at a final concentration of 50 nM. Transfections for the generation of stable cell lines were performed using X-tremeGENE^TM^ 9 DNA transfection reagent (Merck), following the instructions of the manufacturer. Briefly, pcDNA5/FRT/TO and the pOG44 vector (three times the concentration of pcDNA5/FRT/TO plasmid) (Thermo Fischer Scientific) were incubated for 15 min at room temperature with X-tremeGENE^TM^ 9 DNA. The mixture was added to the cells, which were then grown as described above.

Cells were arrested in G1/S phase by treatment with thymidine (Sigma) or in mitosis by treatment with DME (Eg5 inhibitor III, Enzo Life Sciences). For a double thymidine block, cells were first treated with 2 mM thymidine for 16 h, followed by a release from the block for 8 h. The cells were treated again with thymidine for 16 h and released for 8 to 9 h. For flow cytometry analysis, cells were synchronized by a double thymidine block after 6 h of transfection with siRNAs. After the second block, cells were released for 5 h. After addition of 1 µM of DME from a 10 mM stock in DMSO and incubation for 7 h, cells were harvested every 15 min for 120 min from the time of release.

For λ-protein phosphatase (λPP) treatment, asynchronous or mitotic HeLa cell lysates were incubated with 0.9 mM MnCl_2_ and 7 U/µl λPP (New England Biolabs) or 0.9 mM MnCl_2_ and water for 1 h at 30 °C.

### Generation of stable cell lines

HeLa Flp-In TREx cells were seeded onto 24-well plates and cultured for 24 h. Cells were then co-transfected with 0.5 µg of pcDNA5/FRT/TO-plasmid and 1.5 µg of pOG44 using 1 µl of X-tremeGENE^TM^ 9 DNA. On the next day, cells from three wells were pooled, reseeded in 10 cm dishes and selected using hygromycin (200 µg/ml) and blasticidin (5 µg/ml). Transgene expression was induced by the addition of tetracycline (1 µg/ml) and further incubation for 24 h.

### Protein expression and purification

GST-MSP constructs were expressed in *E. coli* BL21 (DE3) codon^+^ strain at 16 °C for 16 h upon induction with 1 mM isopropyl β-D-1-thiogalactopyranoside (IPTG). After washing with PBS, cells were suspended in lysis buffer (50 mM Tris-HCl, pH 7.5, 300 mM NaCl, 1 mM MgCl_2_, 1 mM DTT, 1 mg/ml each of aprotinin, pepstatin and leupeptin and 0.1 mM PMSF) and lysed using an Emulsiflex-C3 (Avestin, Germany). The lysate was centrifuged at 100,000 g at 4 °C for 30 min and the supernatant was incubated using Glutathione Sepharose-4-Fast Flow resin for 1 h at 4 °C followed by washing steps with lysis buffer. Proteins were eluted using lysis buffer supplemented with 15 mM glutathione and dialyzed overnight at 4 °C against lysis buffer.

### Peptide pull down assays

Synthetic biotinylated peptides were purchased from ProteoGenix, France. Peptide pull down assays were performed as described by (Di Mattia et al., 2020). Briefly, for peptide pull down assays from total HeLa-cell lysate, 30 nmol of biotinylated peptides were incubated with 30 µl neutravidin agarose resin in 1 ml of buffer 1 (50 mM Tris– HCl, pH 7.4, 75 mM NaCl, 1 mM EDTA, 1% Triton X-100 and protease inhibitor cocktail tablet (cOmplete, Roche)) at 4 °C for 1 h. The immobilized beads were then washed twice with buffer 2 (50 mM Tris–HCl, pH 7.4, 500 mM NaCl, 1 mM EDTA, 1% Triton X-100 and protease inhibitor cocktail tablet (cOmplete, Roche)) and twice with buffer 1. HeLa P4 cells from one 10 cm dish (8 million cells) per condition were lysed using 500 µl of buffer 3 (50 mM Tris–HCl, pH 7.4, 75 mM NaCl, 1 mM EDTA, 1% Triton X-100, protease inhibitor cocktail tablet (cOmplete, Roche) and phosphatase inhibitor (phosSTOP) tablet (Roche)). The cell lysate was incubated for 5 min on ice and centrifuged at 9,600 g at 4 °C for 10 min. The cleared lysate was incubated with peptide-coupled neutravidin beads at 4 °C for 2 h. The beads were then washed four times with 1 ml buffer 3 and bound proteins were eluted using 30 µl of SDS sample buffer (4% (w/v) SDS, 125 mm Tris, pH 6.8, 10% (v/v) glycerol, 0.02% (v/v) bromophenol blue, and 10% (v/v) β-mercaptoethanol)).

For binding assays using peptides and recombinant proteins, 20 nmol of biotinylated peptides were incubated with 20 µl neutravidin agarose resin in 500 µl of buffer 4 (50 mM Tris–HCl, pH 7.4, 75 mM NaCl, 1 mM EDTA, 1% Triton X-100, 1 mM DTT and protease inhibitor cocktail tablet (cOmplete, Roche)) at 4 °C for 1 h. The immobilized beads were then washed thrice with buffer 4 and blocked using 1.25 mg/ml hemoglobin from bovine blood (Sigma) at 4 °C for 1 h. The beads were washed thrice with buffer 4. Thirty micrograms of purified proteins were incubated in 500 µl buffer 5 (50 mM Tris– HCl, pH 7.4, 75 mM NaCl, 1 mM EDTA, 1% Triton X-100, 0.25 mM DTT and protease inhibitor cocktail tablet (cOmplete, Roche)) at 4 °C for 2 h. The beads were then washed four times with buffer 5 and bound proteins were eluted using 25 µl of SDS sample buffer.

### Immunoprecipitation

2x 10^6^ HeLa Flp-In TREx cells were seeded per 10 cm dish with or without the addition of 1 µg/ml tetracycline. After 24 h, the cells were treated with either 2 mM thymidine or 0.5 µM DME for 14 h to arrest them in G1/S or in mitosis, respectively. The cells were lysed using lysis buffer (10 mM Tris–HCl, pH 7.4, 400 mM NaCl, 2 mM EDTA, 1% Triton X-100, 0.1% SDS, 1 mM DTT, 10 U/ml benzonase (EMD Millipore), protease inhibitor cocktail tablet (cOmplete, Roche) and phosphatase inhibitor (phosSTOP) tablet (Roche)) and then centrifuged for 15 min at 14,000 g at 4 °C. The clarified lysate was then diluted 3.75 times using dilution buffer (10 mM Tris–HCl, pH 7.4, 2 mM EDTA, 1 mM DTT, protease inhibitor cocktail tablet (cOmplete, Roche) and phosphatase inhibitor (phosSTOP) tablet (Roche)). For immunoprecipitation of GFP-VAPB, 50 µl of GFP-Selector (NanoTag) were washed thrice with 3.75 times diluted lysis buffer. The lysate was incubated with the beads for 3 h at 4 °C. The beads were then washed four times with diluted lysis buffer and bound proteins were eluted with 40 µl of SDS sample buffer. For immunoprecipitation of endogenous protein complexes from total HeLa cell lysate, mitotic cells were enriched as described above. 4 μg of rabbit anti-VAPB, or IgG as a control were immobilized on 50 μl of Protein A–Sepharose 4 Fast Flow beads (GE Healthcare) for 3 h and incubated with clarified lysates.

### GST pull down

GST proteins (200 pmol) were immobilized on GST-Selector beads (Nanotag, Germany) equilibrated in diluted lysis buffer as described above (see immunoprecipitation) for 1 h at 4 °C. The beads were washed thrice using diluted lysis buffer. Mitotic HeLa P4 cell lysates (see immunoprecipitation protocol) were then incubated with immobilized beads for 3 h. Bound proteins were eluted with 40 µl of SDS sample buffer.

### Flow cytometry

HeLa P4 cells released from a double thymidine block were fixed with 70% ethanol overnight at 4 °C. The fixed cells were permeabilized with 0.25% Triton X-100 in PBS for 5 min and blocked with 1% BSA for 5 min. The cells were stained with an Alexa Fluor 488-labeled anti-phospho-histone H3 (ser28) antibody (Thermo Fischer Scientific) for 90 min. After staining, cells were resuspended in PBS containing RNase A (Machery-Nagel, Germany) (0.1 mg/ml) and incubated overnight at 4 °C. Propidium iodide (Thermo Fischer Scientific) was added to a concentration of 0.03 µg/µl and cells were analyzed using a BD FACSCanto II (Becton-Dickinson). Data was analyzed with FlowJo software.

### Phosphopeptide enrichment and LC-MS

Protein samples were separated by SDS-PAGE and Coomassie stained bands were cut out. Proteins were reduced, alkylated and digested in-gel with the endoproteinase trypsin. Peptides were extracted and phosphopeptides were enriched by TiO_2_ chromatography as described (Oellerich et al., 2009). In brief, peptides were dissolved with 20 µl of 5% (v/v) glycerol in 80% (v/v) acetonitrile (ACN), 5% (v/v) trifluoroacetic acid and loaded onto a TiO_2_ column. The column was washed 3 times with 20 µl of 5% (v/v) glycerol in 80% (v/v) ACN, 5% (v/v) trifluoroacetic acid and 5 times with 20 µl of 80% (v/v) ACN, 5% (v/v) trifluoroacetic acid. The column was then incubated 3 times with 20 µl of 0.3 normal (N) NH_4_OH, pH ≥ 10.5, to elute phosphopeptides. LC-MS analysis of phosphopeptides was performed under standard conditions on a Lumos Tribrid mass spectrometer (ThermoFisherScientific) working in data-dependent acquisition mode using top 15 method. MS was coupled to an UltiMate LC system (ThermoFisherScientific) equipped with C18 trap column packed in-house (1.5 cm, 360-μm outer diameter, 150-μm inner diameter, Nucleosil 100-5 C18; MACHEREY-NAGEL, GmbH & Co. KG) and an analytical C18 capillary self-made column (30 cm, 360-μm outer diameter, 75 μm-inner diameter, Nucleosil 100-5 C18). The flow rate was 300 nL/min, with a gradient from 8 to 40% (v/v) ACN in 0.1% (v/v) formic acid for 40 min. Data were processed with MaxQuant software (MQ Version 1.6.17.0) and searched against the Uniprot database (taxonomy human, October 2017, 48705 entries) with phosphorylation at serine, threonine, and tyrosine residues and methionine oxidation as variable modifications, and cysteine carboxyamidomethylation as fixed modification. The precursor tolerance was set to 10 ppm and the MS/MS tolerance to 20 ppm. Data were reported with 1 % FDR.

### Western blot analyses

Western blotting was performed according to standard protocols using IRDye (LI-COR) secondary antibodies. 3-8% NuPAGE Tris-Acetate gels (Thermo Fischer Scientific) were used to separate ELYS by SDS-PAGE. For other proteins, 4-12% NuPAGE Bis-Tris gels were used. After transfer to PVDF or nitrocellulose, the membranes were incubated in blocking buffer (3% BSA in TBS-T (24.8 mM Tris, pH 7.4, 137 mM NaCl, 2.7 mM KCl, 0.05% (v/v) Tween 20)) for 1 h at room temperature and with primary antibodies (Table 2) overnight at 4 °C. Incubation with IRDye (diluted 1:10,000 in blocking buffer) for 1 h at room temperature was followed by three washing steps with TBS-T. Proteins were detected using LI-COR Odyssey-CLx imaging system and analyzed by Image Studio Lite software (LI-COR). For statistical analysis, ordinary one-way Anova, followed by Brown-Forsythe and Welch tests were performed using GraphPad Prism 8 software and a confidence interval of 95% was set.

**Table 2:**
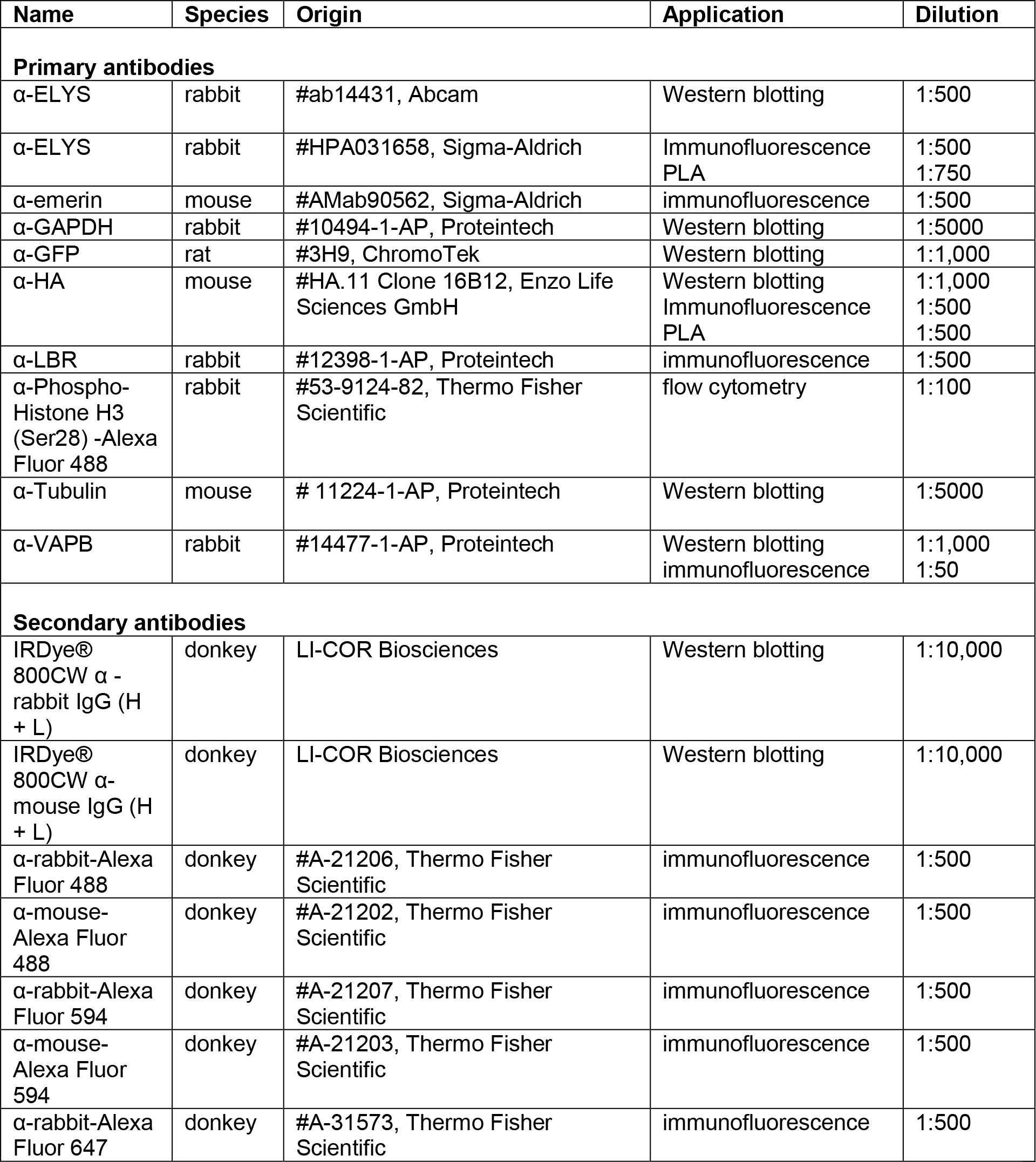
Antibodies used in this study.

| Name | Species | Origin | Application | Dilution |
| --- | --- | --- | --- | --- |
| **Primary antibodies** |  |  |  |  |
| α-ELYS | rabbit | #ab14431, Abcam | Western blotting | 1:500 |
| α-ELYS | rabbit | #HPA031658, Sigma-Aldrich | Immunofluorescence<br>PLA | 1:500<br>1:750 |
| α-emerin | mouse | #AMab90562, Sigma-Aldrich | immunofluorescence | 1:500 |
| α-GAPDH | rabbit | #10494-1-AP, Proteintech | Western blotting | 1:5000 |
| α-GFP | rat | #3H9, ChromoTek | Western blotting | 1:1,000 |
| α-HA | mouse | #HA.11 Clone 16B12, Enzo Life Sciences GmbH | Western blotting<br>Immunofluorescence<br>PLA | 1:1,000<br>1:500<br>1:500 |
| α-LBR | rabbit | #12398-1-AP, Proteintech | immunofluorescence | 1:500 |
| α-Phospho-Histone H3 (Ser28) -Alexa Fluor 488 | rabbit | #53-9124-82, Thermo Fisher Scientific | flow cytometry | 1:100 |
| α-Tubulin | mouse | # 11224-1-AP, Proteintech | Western blotting | 1:5000 |
| α-VAPB | rabbit | #14477-1-AP, Proteintech | Western blotting<br>immunofluorescence | 1:1,000<br>1:50 |
| **Secondary antibodies** |  |  |  |  |
| IRDye® 800CW α - rabbit IgG (H + L) | donkey | LI-COR Biosciences | Western blotting | 1:10,000 |
| IRDye® 800CW α- mouse IgG (H + L) | donkey | LI-COR Biosciences | Western blotting | 1:10,000 |
| α-rabbit-Alexa Fluor 488 | donkey | #A-21206, Thermo Fisher Scientific | immunofluorescence | 1:500 |
| α-mouse-Alexa Fluor 488 | donkey | #A-21202, Thermo Fisher Scientific | immunofluorescence | 1:500 |
| α-rabbit-Alexa Fluor 594 | donkey | #A-21207, Thermo Fisher Scientific | immunofluorescence | 1:500 |
| α-mouse-Alexa Fluor 594 | donkey | #A-21203, Thermo Fisher Scientific | immunofluorescence | 1:500 |
| α-rabbit-Alexa Fluor 647 | donkey | #A-31573, Thermo Fisher Scientific | immunofluorescence | 1:500 |

### Immunofluorescence, confocal microscopy and live cell imaging

For confocal microscopy, cells were grown on coverslips and fixed with 4% (v/v) paraformaldehyde. For immunofluorescence, fixed cells were permeabilized with 0.5% (v/v) Triton X-100 in PBS for 5 min at room temperature and blocked with 3% (w/v) BSA in PBS for 20 min at room temperature. Staining was performed for 1 h at room temperature using appropriate primary antibodies and fluorescently labeled secondary antibodies (Table 2), which were diluted in 3% BSA in PBS. Cells were embedded in MOWIOL-supplemented with 1 µg ml^-1^ DAPI. Cells expressing fluorescently labeled proteins were mounted directly with MOWIOL containing DAPI.

An LSM510 confocal laser scanning microscope using a 100X /1.3 oil immersion lens (Zeiss, Germany) was used for microscopic analysis. For live cell imaging, cells were grown on ibidi µ-Dish^35^ ^mm, low^ and imaged using an LSM800 confocal laser scanning microscope using a 40X/1.3 oil DIC M27 immersion lens (Zeiss) under 5% CO_2_ at 37 °C. Images were analyzed using Zen Blue software (Zeiss). For statistical analysis, ordinary one-way Anova followed by Brown-Forsythe and Welch tests was performed and a confidence interval of 95% was set.

### Proximity Ligation Assay (PLA)

HeLa Flp-In TREx cells expressing HA-VAPB or HA-VAPB-KD/MD were seeded at a density of 35,000 cells/well in 24-well plates. After synchronizing the cells in mitosis, cells were fixed with 4% paraformaldehyde and permeabilized with 0.5% (v/v) Triton X-100. Duolink In Situ PLA kit (Sigma Aldrich) was used for PLA. Cells were blocked and incubated with mouse anti-HA or rabbit anti-ELYS antibodies and thereafter with the corresponding PLA probes. After ligation and amplification using the kit reagents, cells were counterstained for HA and ELYS and mounted using Duolink mounting medium with DAPI. Images were acquired on a LSM510 confocal laser scanning microscope using 100X /1.3 oil immersion lens. Anaphase cells were analyzed for PLA interaction using CellProfiler (Carpenter et al., 2006). A pipeline was generated to measure PLA signal intensities and number around the chromatin. Chromatin was identified first by DAPI staining. The region around the chromatin was defined by the function ExpandOrShrinkObjects with a pixel number of 4 for expansion. An unpaired t-test was performed for statistical analysis using GraphPad Prism 8 software and a confidence interval of 95% was set.

## Acknowledgement

We would like to thank Dr. Thomas Mayer (University of Konstanz, Germany) for stable cell lines, Dr. Holger Bastians, (University of Göttingen, Germany) for very helpful advice and reagents and Dr. Larry Gerace, La Jolla, USA for very fruitful discussions. The project was supported by the Deutsche Forschungsgemeinschaft (DFG, SFB1190, P07).

## Supporting Information

**Video 1**

Mitosis in mCherry-H2B HeLa Flp-In TREx cells treated with non-targeting siRNA

**Video 2**

Mitosis in mCherry-H2B HeLa Flp-In TREx cells treated with siVAPB

## Abbreviations

ELYS: embryonic large molecule derived from yolk sac
FFAT: two phenylalanines in an acidic tract
INM: inner nuclear membrane
LAP: lamina associated polypeptide
LBR: lamin B receptor
MSP: major sperm protein<colcnt=2>
NE: nuclear envelope
NPC: nuclear pore complex
ONM: outer nuclear membrane
PLA: proximity ligation assay
RAPIDS: Rapamycin and APEX dependent Identification of proteins by SILAC
VAPB: vesicle-associated membrane protein-associated protein B

